# crossing3dforest: an R package for evaluating empty space structure in forest ecosystems

**DOI:** 10.1101/2023.02.01.526548

**Authors:** Nicola Puletti, Rossella Castronuovo, Carlotta Ferrara

## Abstract

Traditionally, forest structure is mostly described by vegetative elements; however, the complementary empty space also contributes to the forest spatial structure.
We developed an R package (*crossing3dforest*) to support the entire processing of Terrestrial Laser Scanning point clouds to quantify the size, shape, and connectivity of empty spaces within the mid and low strata of forest stands, using an approach based on the percolation theory. The package functions, which are designed for step-by-step single stand analysis, can be executed sequentially in a pipeline.
A case study is presented to demonstrate the *crossing3dforest* potentials for characterising the forest empty space architecture. TLS point clouds collected in ten different pure beech (Fagus sylvatica L.) stands, representative of five distinct forest management regimes, were analysed and characterised.
The adopted empty space approach can be integrated into forest structural analysis to identify animal-habitat associations and establish appropriate habitat structure for wildlife management.

**Graphical abstract:** 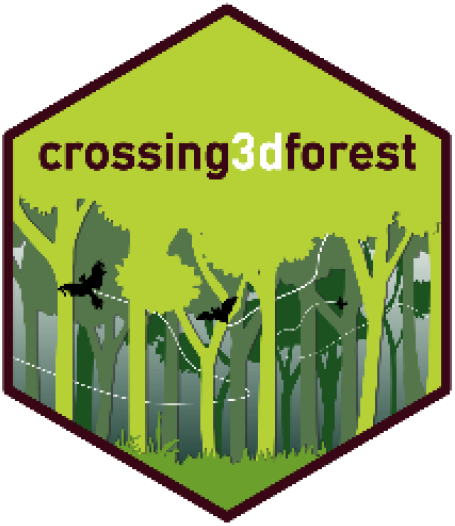

## 1. Introduction

Forest structure, defined as the composition, organisation, and location of above-ground vegetative structures, is a crucial component of ecosystems at global level (Migliavacca et al., 2021). Studies on forest structure have traditionally focused primarily on the vegetative elements and their architecture (Chianucci et al., 2014), but the complementary empty space can also be used to characterise forest structure (MacArthur & Horn, 1969; Lefsky et al., 1999). The “forest air space” (Su et al., 2019) can be defined as a spatial matrix formed between tree stems, branches, and leaves. This space is an essential aspect of forest ecology for the development and evolution of the forest environment. One of the key functions of the empty space is in regulating local air temperature. Recent studies have shown that the empty spaces within a forest can have a significant impact on microclimatic conditions, regulating local air temperature (Zellweger et al., 2020). Furthermore, the amount of available empty space beneath the dominant forest canopy cover serves as an environmental resource for the life cycle of insects (Müller & Brandl, 2009), birds), and micromammals (Blakey et al., 2017). These traits, together with disturbance regimes, global climate, and forest management, influence the different spatial arrangements of both canopy and under-the-canopy spaces, thereby determining the variety of forest stands structures(Decuyper et al., 2018).

Up to now, only a few field methods based on the installation of complex tools have been used for characterising empty space in forest environments (Dial et al., 2004). These methods are time consuming and can be actually limited to small areas. Terrestrial Laser Scanning (TLS) opens a new avenue for a high-accuracy representation of the three-dimensional forest structure from a ground-based perspective up to the canopy level (Calders et al., 2020). TLS surveys are often faster and more accurate than conventional ones, having the possibility of easily scanning multiple locations, even in dense forest stands.

TLS point clouds derived from single or multiple laser scanning approaches can be used to quantify unique high-resolution forest structure features through a variety of indicators (Campos et al., 2021) such as tree volume (Calders et al., 2015 ; Giannetti et al., 2020) above-ground biomass (Puletti et al., 2020), single tree structure (Juchheim et al., 2017), tree-crown (Chianucci et al., 2020), and stand-level characteristics (Grotti et al., 2020; Puletti et al., 2021b)

The *crossing3dforest* package, based on R statistical software, is specifically developed to process and analyse TLS point clouds in order to assess specific information on the size, shape and connectivity of the 3D empty space within the forest stand, particularly under the dominant forest canopy. The underlying approach adopted by this package to assess the empty space features relies on percolation theory. Percolation theory is a branch of physics that deals with the study of the behaviour of a system of connected components undergoing random occupancy. It models the spread of a substance, such as water, gas or electricity, through a porous medium or a network of connected sites. The theory aims to determine the critical probability at which the system switches from being disconnected to becoming connected. Percolation theory has a wide range of applications, including the study of transport phenomena in porous materials, computer networks, and social networks. It also plays a role in understanding the behaviour of complex systems, such as materials with random internal structure and random graphs, and traditionally is used to describe the properties of an object related to the connectivity of its constituent parts (Stauffer & Aharony, 2018;Hunt et al., 2014). For example, it has been widely used in geology to examine how fluid flow in porous media, as well as in landscape ecology for a variety of applications mainly oriented to evaluate landscape patterns and analyse ecological limitations for wildlife traditionally adopting a 2D raster-approach. The most relevant characteristic, as with graph and network theories (Farage et al., 2021) to which percolation theory is related, is the ability in providing overall properties from local, detailed, specifications of the object of interest.

In this paper, we present an applicative example of the empty-space structure being assessed in ten different European Beech (*Fagus sylvatica* L.) forest stands representing five different management regimes: managed coppices (C); unmanaged coppices (UC); high forests (HF); mature forests (MF); old-growth forests (OGF).

## 2. Package overview

*crossing3dforest* has been developed as an R package (R Core Team, 2022). Currently, the most up-to-date version of the package can be downloaded free of charge from the GitLab development repository of the project (https://gitlab.com/Puletti/crossing3dforest).

The key steps to process TLS data using the *crossing3dforest* package are basically:

1. import a normalized point cloud;
2. point cloud voxelization (Puletti et al., 2021b);
3. percolating cluster identification and measuring.

To address the above-mentioned features, the key R package dependencies are:

a. *lidR* (https://cran.r-project.org/web/packages/lidR/index.html) which ensures fast import of raw point cloud files, both las or .laz file format (see § 2.1);
b. *mmand* (https://cran.rstudio.com/web/packages/mmand/index.html) which allows finding connections between voxels. The function *percolation_sts* is based on the function *components*, imported from this R package.

The processing workflow is depicted in Figure 1, with further detail provided in the following sections and summarised in Table 1.

**Table 1.** Main functions included in the package.

| Function (arguments) | Description |
| --- | --- |
| **VoxFor**(inLas,<br>minXrect, maxXrect, minYrect, maxYrect,<br>x.vox, y.vox, z.vox,<br>min.npXvox, Zcut) | Voxelization and classification of voxels in “empty space” and “vegetation” |
| **df2array**(voxgrid.plot, z.min, z.max) | Creates the input for percolation_sts function |
| **percolation_sts**(x, k) | Defines connections between voxels using a 'von Neumann 3d connections' kernel |
| **plotVoxel**(xc,yc, zc,<br>x.vox, y.vox, z.vox,<br>vox.col, alpha) | Add voxels to a rgl 3-dimensional graph |

**Figure 1.**
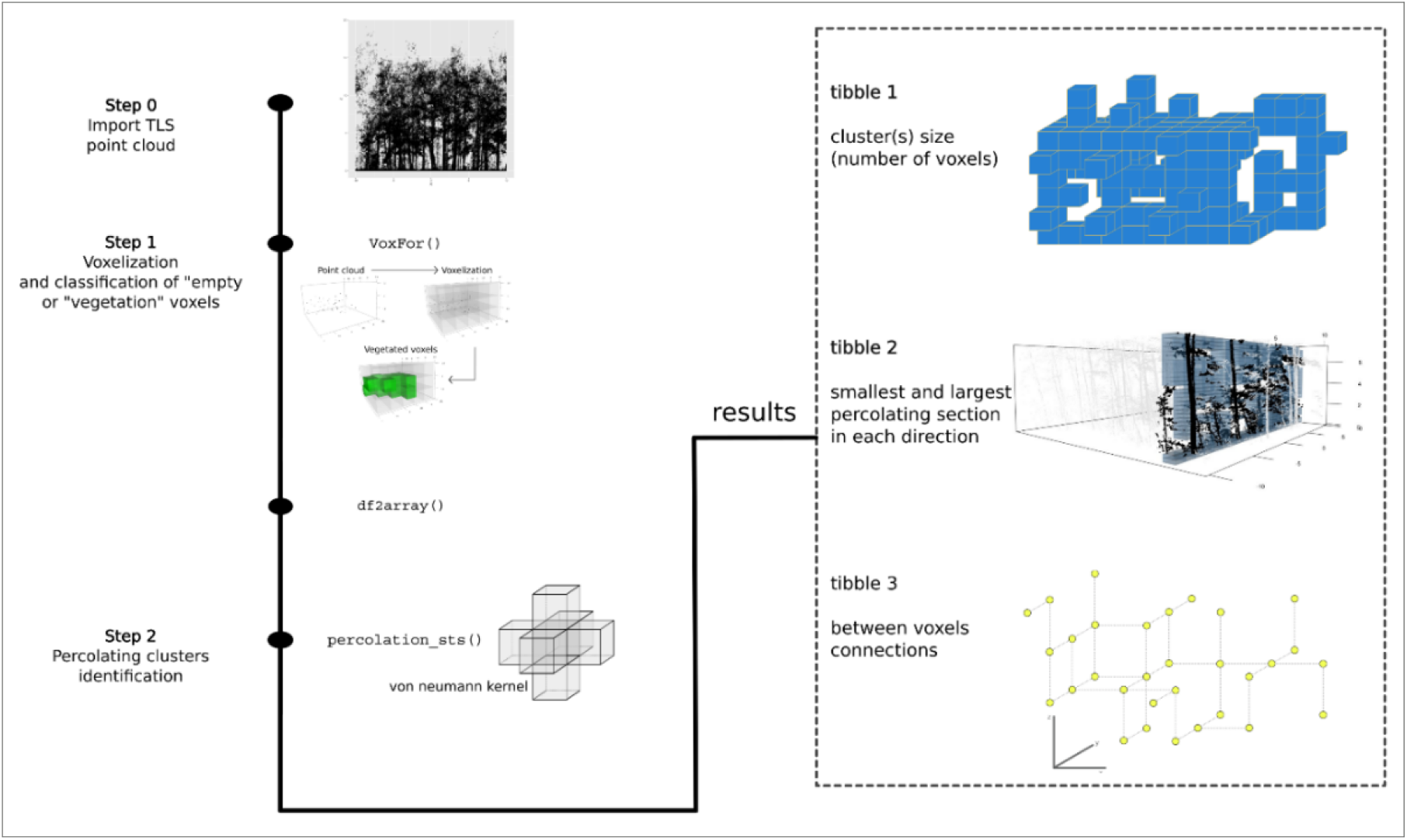
*crossing3dforest* workflow steps and functions for deriving empty space parameter characterization.

### 2.1. Import TLS data

Point clouds collected in forest environments require a first step of “point cloud normalisation”. The *crossing3dforest* package requires already normalised point clouds in .las or .laz format that can be managed using the functionalities of *lidR* package. Such *lidR* object is the first argument of the *VoxFor* function.

### 2.2. Voxel grid definition

The second step of the process is twofold. First, a user defined voxel grid is created, then the number of points within each voxel is calculated. These two phases are particularly important in optimising the computational process. Using the specific arguments of the *VoxFor* function, the user is able to crop the entire area of interest and define the voxel sizes in the x, y and z dimensions.

*VoxFor* function computes the number of points within each voxel of the user-defined 3D-grid. Each voxel is then classified as either “vegetated” or “empty space” depending on the threshold parameter *min.npXvox*: voxels comprising a number of point higher than *min.npXvox* will be classified as “vegetation” otherwise they are considered “empty space”.

The output of this function is a *data.frame* object containing (1) the coordinates of the voxels (minimum, maximum and central coordinates), (2) the number of points in each voxel, and (3) their classification (1 = empty voxel; 0 = vegetation voxel).

### 2.3. Create the array

The *df2array* function requires the *data.frame* obtained from *VoxFor* function (par 2.2) and the minimum and maximum values of the Z layer of interest. The output is a list of two elements. The first element of the list is the array that will be used in *percolation_sts* function (see par 2.4). The second element can be used for graphical functions (see par. 2.5).

### 2.4. Retrieve percolation metrics

The *percolation_sts* function provides specific information on the amount and shape of empty space within forest stands using structural information collected using TLS. The function defines connections between voxels using the “von Neumann” 3d connections kernel, a computer science term referring to the concept of creating a three-dimensional matrix of interconnected nodes in a computer system. In percolation theory, the “von Neumann” kernel refers to a specific type of lattice structure used to model percolation phenomena, such as the flow of liquids or gases through a porous medium. In this context, the “von Neumann” kernel is a set of connections between lattice sites that defines the topological structure of the lattice and determines the flow patterns in the system.

After voxelization, the function codes with “1” the empty cells and then computes the percolation in each of the 3 dimensions, identifying percolating clusters (in that dimension) and their smallest section (orthogonal to that direction). Moreover, total number of cells is computed for each percolating cluster, in any direction.

### 2.5. Graphical function

The *plotVoxel* function enables the visualisation of classified voxels (“empty” or “vegetation”) in a *rgl* plot. Graphical function parameters can be profitably obtained using the values obtained from functions described above. Figure 2 depicts an example achieved using the code reported in Box S1.

**Figure 2:**
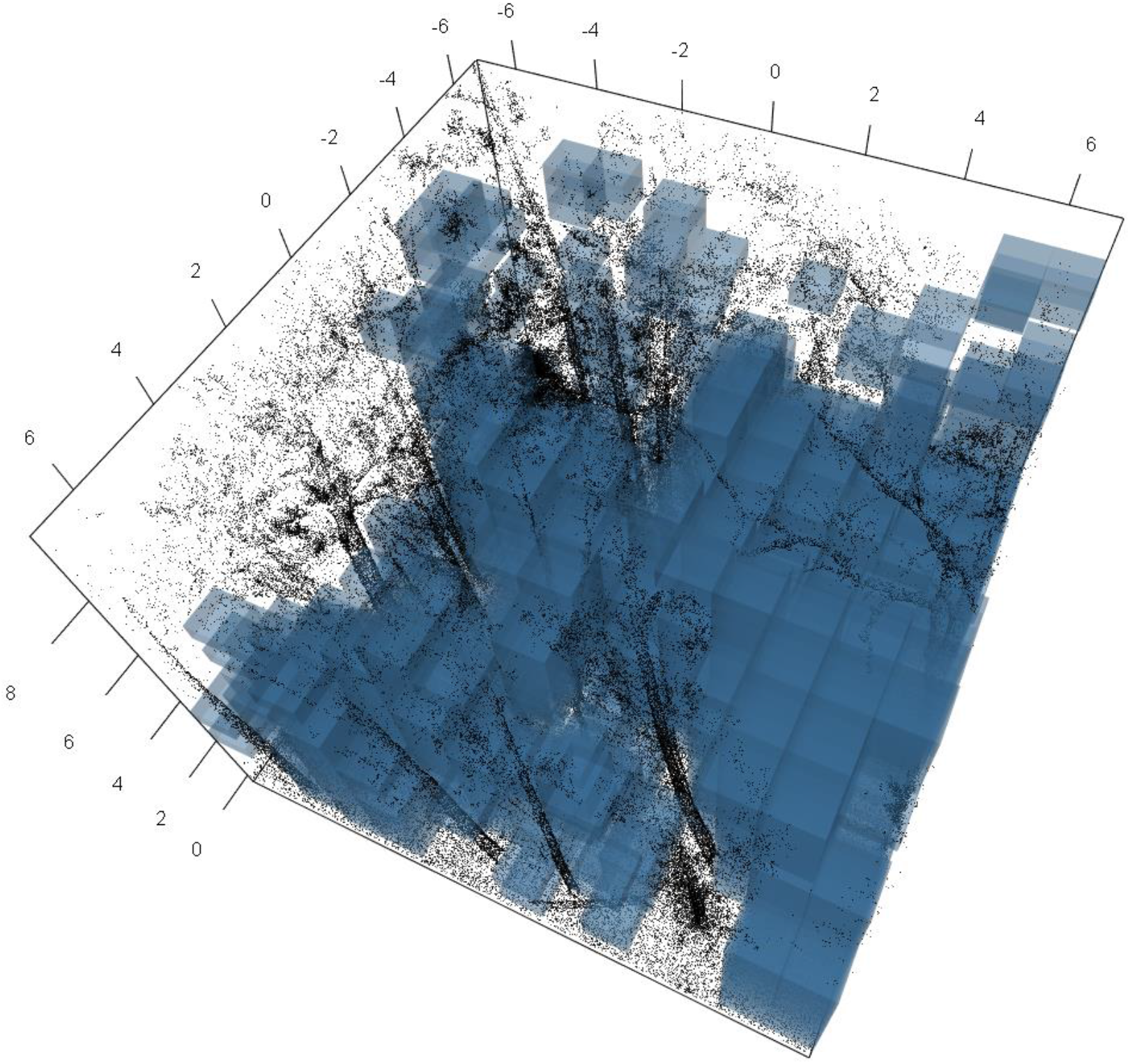
*an example obtained as a screenshot (*rgl.snapshot(’3dplot.png’, fmt = ’png’) *) from the code contained in Box S1*.

## 3. Example application

### 3.1 Stands description, data collection and preprocessing

The examples presented in the following paragraphs aim to demonstrate the ability of *crossing3dforest* package in deriving new forest structural indicators. We used data collected in ten stands dominated by European Beech (Fagus sylvatica L., 1753) under five different forest management regimes. The stands show differences in forest structure, as a result of different silvicultural management and age (Table 2). One of the stands (UC1, Figure 3) is located in the Sila National Park, Italy (11°43’37.17”N, 76°27’44.19”E), while the other nine are located in the Foreste Casentinesi Monte Falterona e Campigna National Park (Figure 3).

**Table 2:** Summary characteristics of the considered forest stands including, the years from the last management activity, stem density, quadratic mean diameter at breast height (QMD), basal area (BA), mean total tree height (Htot). Forest mensuration’s were automatically obtained by functions of the TreeLS package available on GitHub (github.com/tiagodc/TreeLS).

| Plot id | Years from last activity (yrs) | Stem Density (N ha<sup>-1</sup>) | QMD (cm) | BA (m<sup>2</sup> ha<sup>-1</sup>) | Htot (m) |
| --- | --- | --- | --- | --- | --- |
| C1 | <5 | 562 | 48 | 37 | 11.5 |
| C2 | <5 | 781 | 55 | 45 | 11.9 |
| UC1 | ~30 | 863 | 18.5 | 25 | 14.8 |
| UC2 | ~50 | 594 | 51 | 47 | 25.0 |
| HF1 | <5 | 261 | 33 | 28 | 29.9 |
| HF2 | ~50 | 225 | 34 | 30 | 26.7 |
| MF1 | >100 | 259 | 49 | 45 | 32.5 |
| MF2 | >100 | 223 | 31 | 27 | 30.0 |
| OGF1 | >150 | 247 | 63 | 53 | 32.9 |
| OGF2 | >150 | 243 | 41 | 35 | 27.0 |

**Figure 3:**
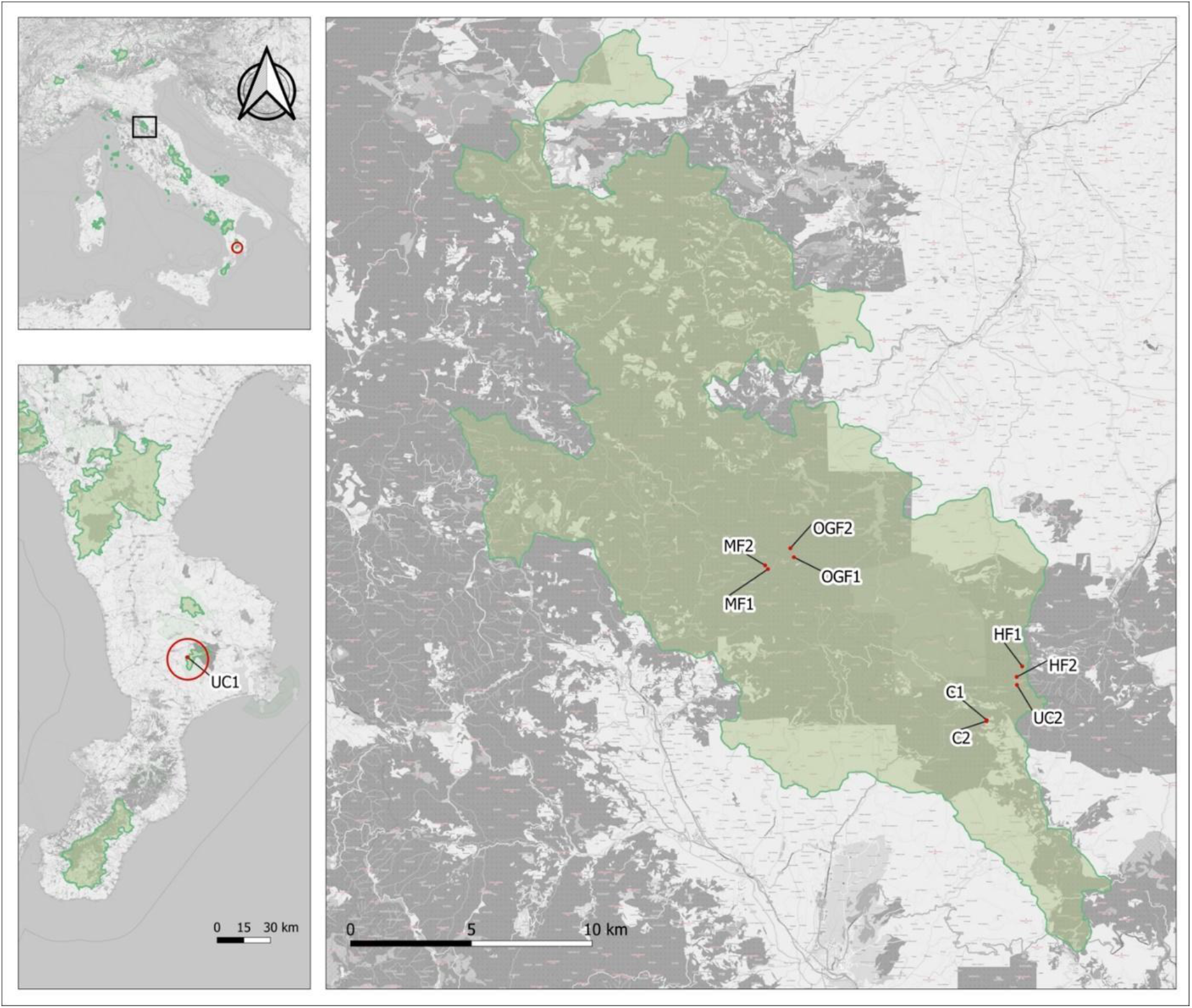
Study areas located in central (black square) and south Italy (red circle), comprising a total of nine plots. One forest stand is located in the Sila National Park (bottom left inset image), while the other nine in the Foreste Casentinesi Monte Falterona e Campigna National Park (right inset image). In green are the boundaries of the National Parks in Italy.

TLS data were collected using FARO® Focus 3D (FARO Technologies Inc.). Following a multiscan approach, 8-12 spherical targets (14 cm diameter) were systematically placed in each plot to allow subsequent co-registration of each individual point cloud (Wang et al., 2014). After point cloud pre-processing steps (Pascu et al., 2019), inner subsets of 20 m x 20 m were extracted from the ten TLS point clouds.

### 3.2. Voxelization

In this experience, a voxel size of .5 x .5 x .25 metres was used (Box 1), with a threshold to define a voxel as “empty” equal to 10 (*min.npXvox* = 10), and the height of maximum point density in the canopy set to 0 (Zcut = 0, i.e. no filter).

**Box 1:**
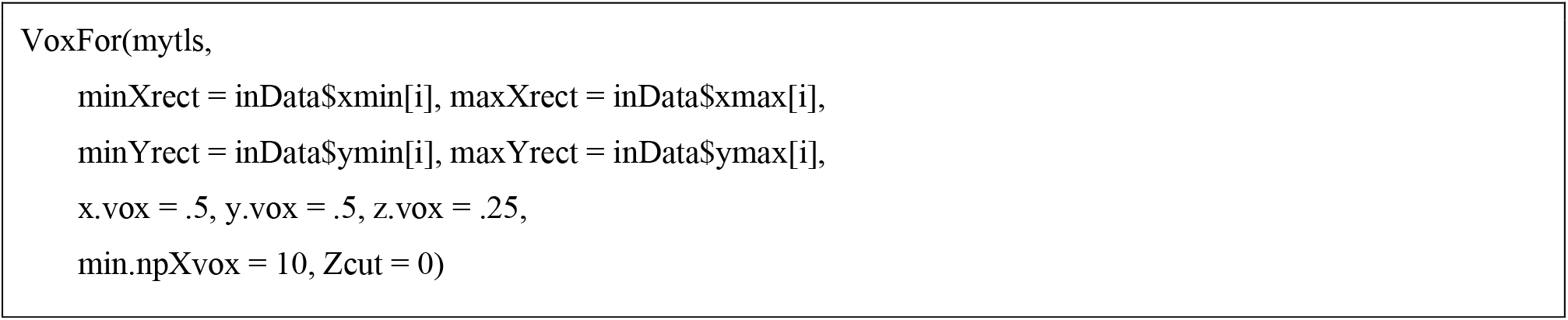
an example of voxelization using the VoxFor() function.

### 3.3. Percolation statistics between stands

The code to compute percolation statistics is presented in Box 2. Table 3 summarises the principal percolation statistics for each forest stand and over the five considered forest management regimes, obtained by *percolation_sts* function.

**Table 3.** Summary of the percolation statistics computed through the crossing3dforest percolation_sts function for each forest stand. (*) Statistic obtained from the 1^st^; (**) statistic obtained from the 2^nd^ tibble.

| plot id | Total number of voxels (# voxel) | Largest cluster size * (#voxel) | Large cluster ratio | Minimum percolating section ** (m<sup>2</sup>) |
| --- | --- | --- | --- | --- |
| C1 | 103,010 | 36,369 | 0.35 | 617.0 |
| C2 | 104,982 | 34,231 | 0.33 | 554.0 |
| UC1 | 125,983 | 44,413 | 0.35 | 736.5 |
| UC2 | 191,984 | 85,401 | 0.44 | 1678.0 |
| HF1 | 228,903 | 131,297 | 0.57 | 2789.0 |
| HF2 | 200,222 | 129,967 | 0.65 | 2684.5 |
| MF1 | 253,830 | 146,702 | 0.58 | 3233.5 |
| MF2 | 230,272 | 133,772 | 0.58 | 2731.5 |
| OGF1 | 242,128 | 122,875 | 0.51 | 2525.0 |
| OGF2 | 221,947 | 110,480 | 0.50 | 2365.5 |

The total number of voxels obviously increases in proportion to the average height of the stand; then, coppice (C) has a lower total number of voxels than mature (MF) or old growth forests (OGF) (Table 3). The largest cluster size statistic (i.e., the size of the largest cluster of connected empty voxels) and its ratio over the total number of voxels (similar to a normalisation), contribute to characterising the forest structure under the canopy.

**Box 3:**
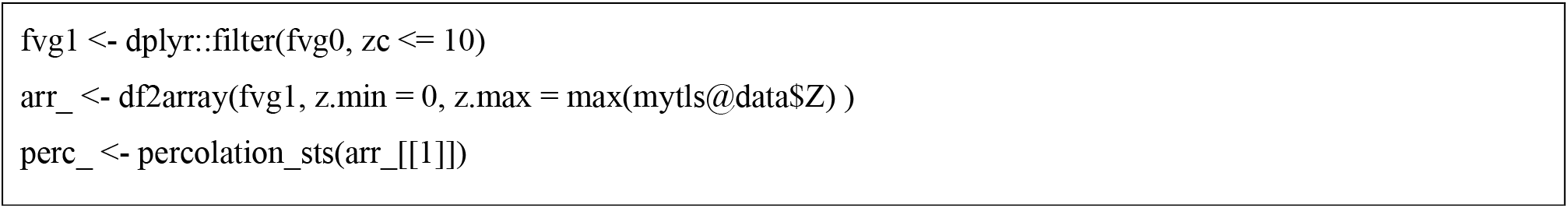
an example for statistical computations.

The output “minimum percolating section” highlights the effects of management regimes in forest structure. In particular, managed coppice stands (C) are characterised by the smallest mean size of percolating sections (585.5 voxels), which can be related to an empty space matrix with a high coefficient of friction. This value slightly increases in unmanaged coppice (UC). From a structural point of view UC stands have a few large trees mixed with a lot of shoots in the understory layer. On the contrary, given the presence of larger empty spaces in mature forest structures, the size of minimum percolating sections increases significantly from coppice to high forest stands.

The same structural dynamic is highlighted by the indicator “number of links” (Figure 4 and 5). In coppice forests, for example, the percentage of highly connected voxels (*n.links* >=5) is less than half of the other management regimes.

**Figure 4.**
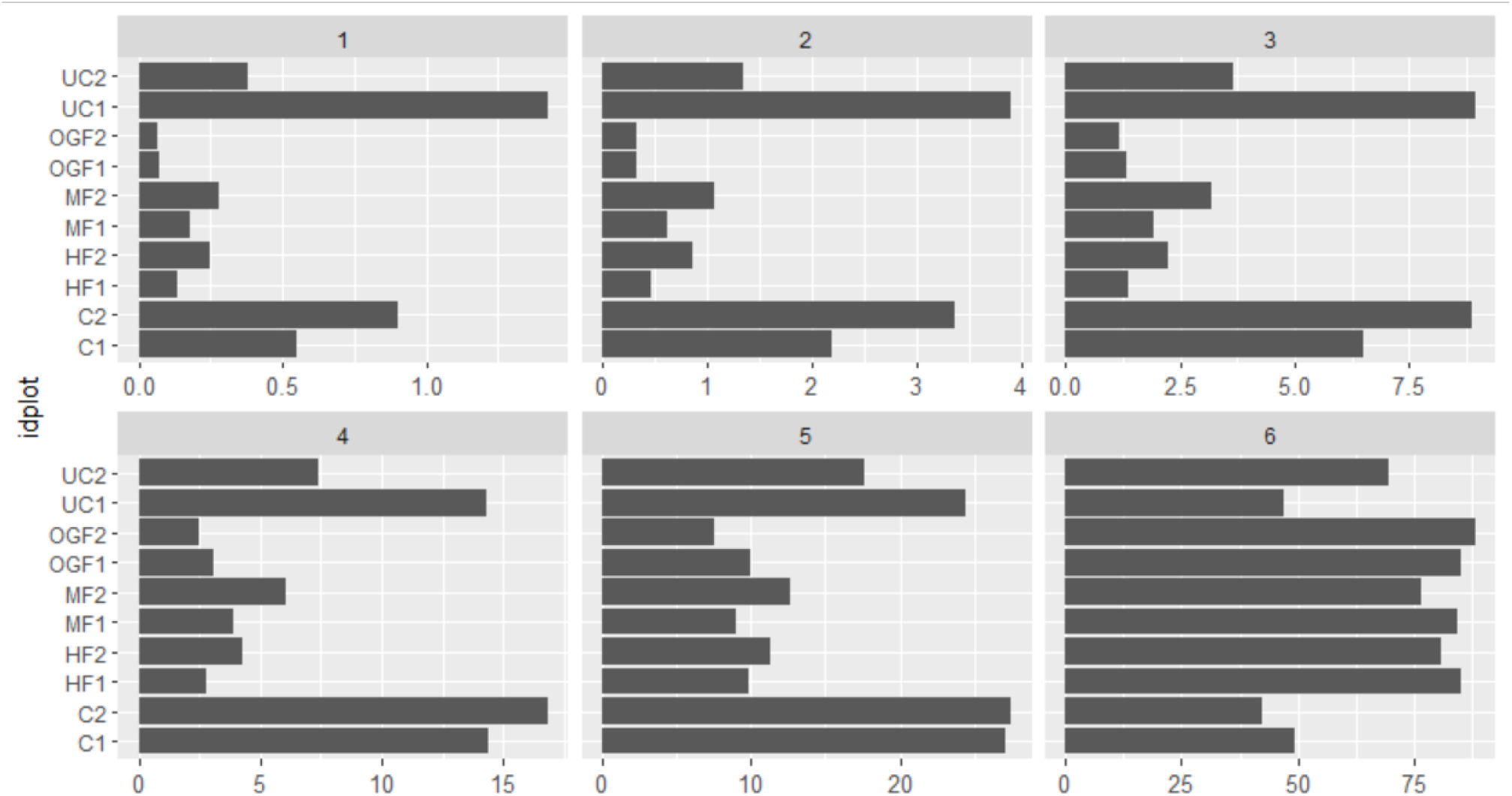
Percentage of the number of links in each forest. This statistic can be derived using the 3^rd^ tibble of the percolation_sts function.

**Figure 5.**
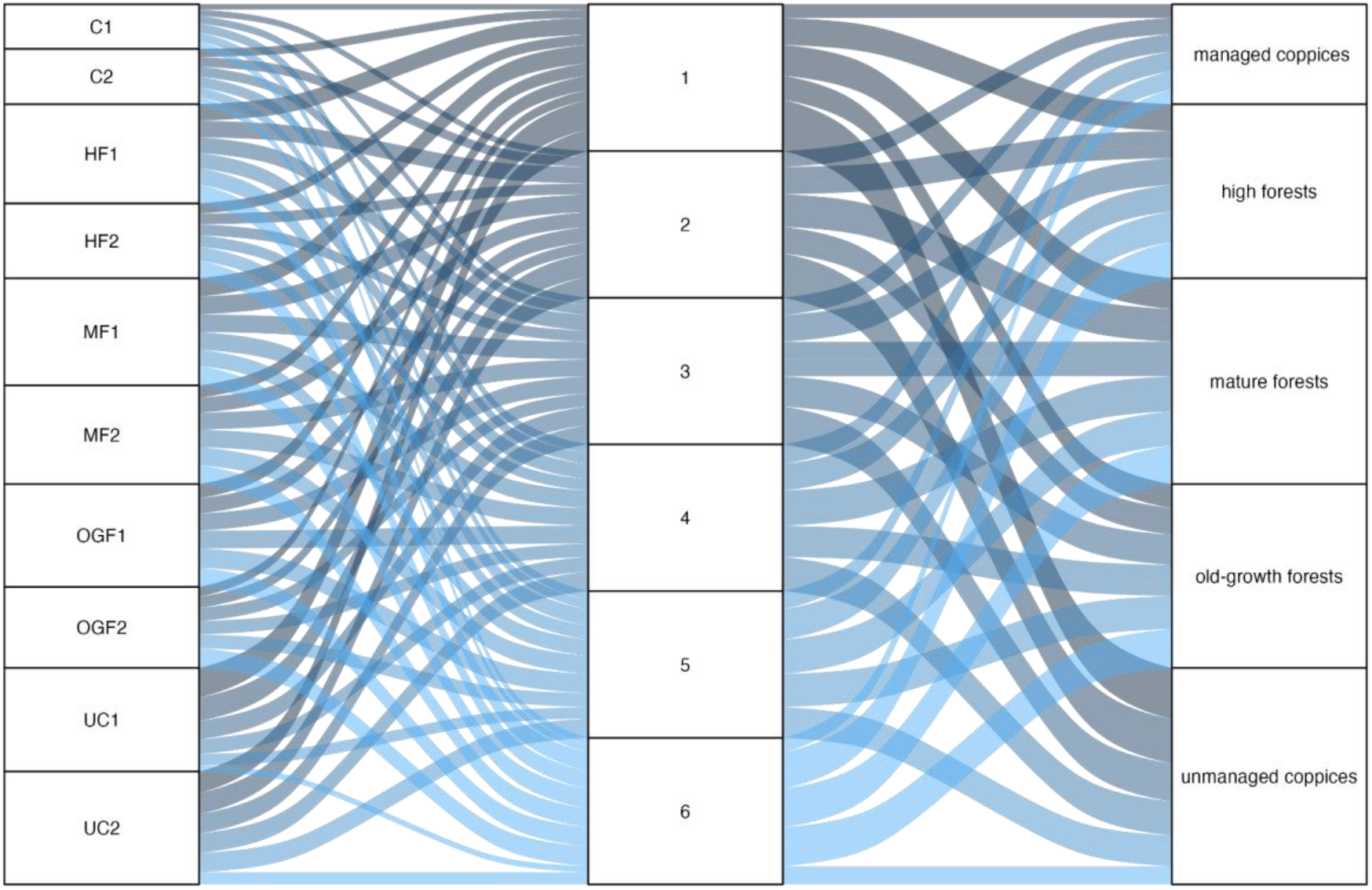
Alluvial diagram illustrating the share of empty space among forest plots and management types. Colors correspond to the number of links as they move through forest stands and management types, with the flow width being proportional to the share percentage. The dimension of rectangles is proportional to the data’ prevalence.

When quantifying percolating empty space with *percolation_sts*, the provided statistics allow to distinguish the principal stand structural traits, which are strictly related to different forest management regimes. In this regard, a hierarchical cluster analysis revealed that 1) coppice management types are clearly distinct from the others, 2) within the coppice cluster, the two managed coppice stands are well separated from the others, 3) among the high forest cluster, the old-growth forests are grouped separately from the others (Figure 6).

**Figure 6.**
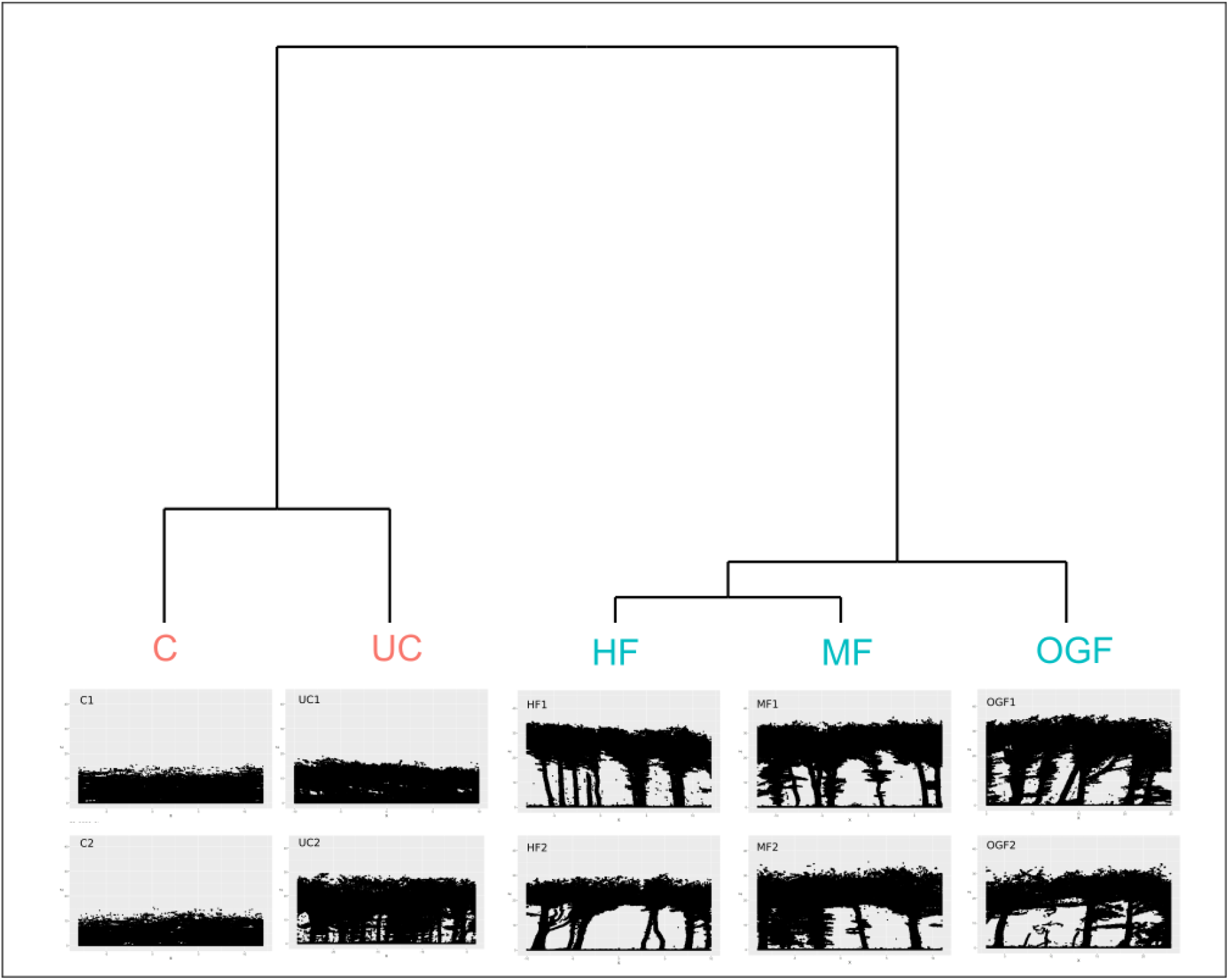
Hierarchical cluster analysis results based on the percolation statistics. On the top: Cluster dendrogram plot relating the management types and forest plots. On the bottom: point cloud sections of the forest plots identified through the cluster analysis.

## 4 Final remarks

The assessment of the attributes beneath the forest canopy is becoming increasingly important in understanding forest ecosystem processes and services, but it is challenging to capture in traditional forest surveys (Ferrara et al., 2023). Although conventional measurements are essential for pre-standardized management decisions, TLS nowadays can significantly improve quantitative monitoring issues in forest environments. *crossing3dforest* package provides tools for implementing a percolation approach to create new structural indicators on the empty space under the canopy.

Further studies should focus on exploring the applications of indicators produced by *crossing3dforest* in different contexts, particularly those related to:

- solve analytical problems (as outlined in the application presented in this work);
- develop indicators for forest management (Berenguer et al., 2014; Decuyper et al., 2018) and for forest fire prevention (Andersen et al., 2004 ; Erdody & Moskal, 2010);
- assess the influence of habitat spatial architecture on animal biodiversity and behaviour (Lecigne et al., 2020);
- analyse the effects of forests on regulating microclimatic conditions (Lindenmayer et al., 2022; De Lombaerde et al., 2021; Su et al., 2019).

## Supporting information

supp mat box 1

## Acknowledgments or Funding?

This research was funded by the Italian Ministry of Agriculture, Food, and Forestry Policies (MiPAAF), subproject “Precision Forestry” of the project AgriDigit program-DM, funding number 36503.7305.2018 of 2 December 2018). A special thanks go to Matteo Guasti (CREA), Simone Innocenti (CREA) and Mirko Grotti (Università La Sapienza) for their valuable support in field data collection.

---

**Box S1:**
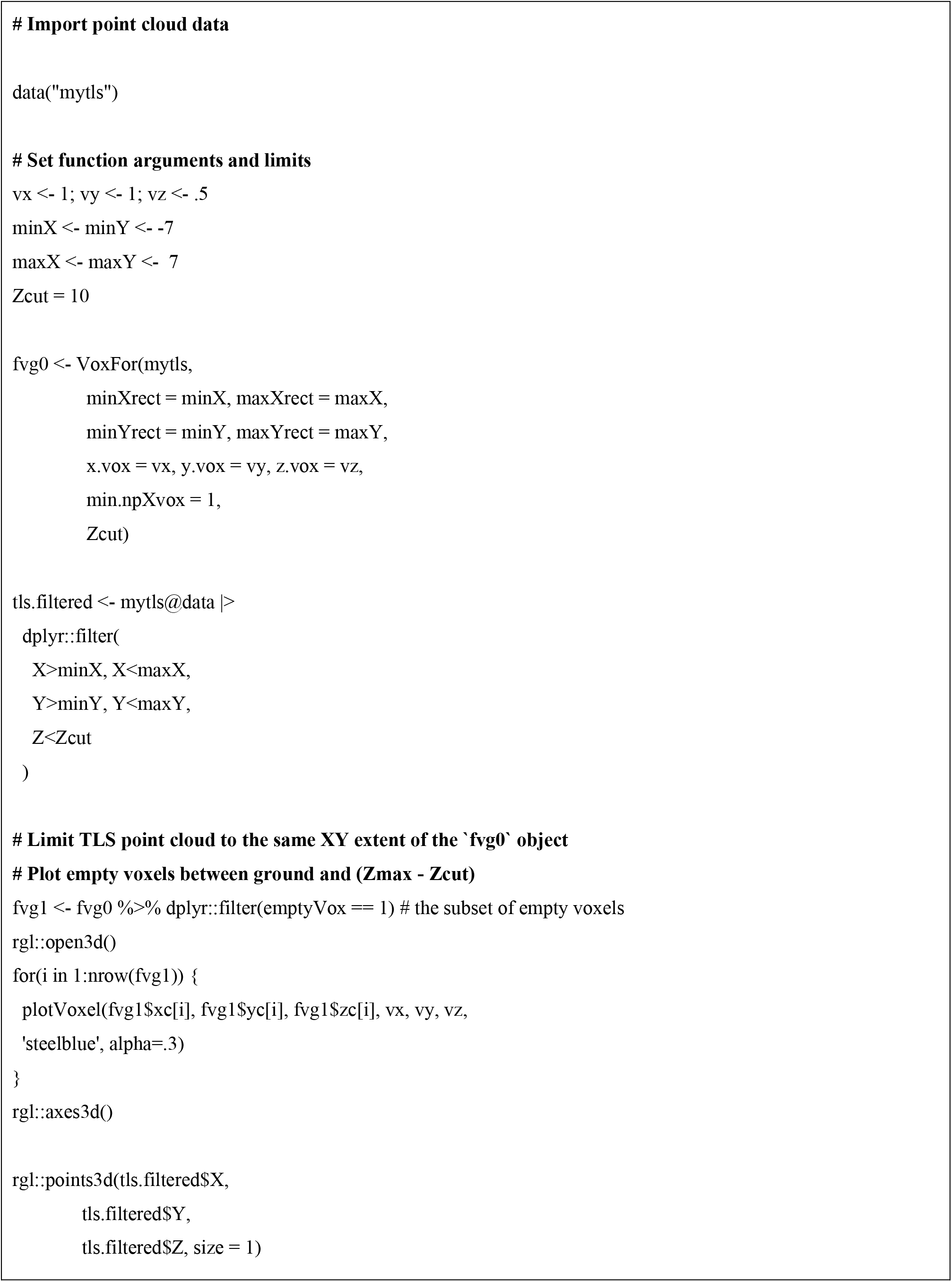
code used to produce figure 2.

