## Supplementary material for "crossing3dforest: an R package for evaluating empty space structure in forest ecosystems": supp mat box 1

```
# Import point cloud data

data("mytls")

# Set function arguments and limits
vx <- 1; vy <- 1; vz <- .5
minX <- minY <- -7
maxX <- maxY <-  7
Zcut = 10

fvg0 <- VoxFor(mytls,
        minXrect = minX, maxXrect = maxX,
        minYrect = minY, maxYrect = maxY,
        x.vox = vx, y.vox = vy, z.vox = vz,
        min.npXvox = 1,
        Zcut)

tls.filtered <- mytls@data |>
  dplyr::filter(
    X>minX, X<maxX,
    Y>minY, Y<maxY,
    Z<Zcut
  )

# Limit TLS point cloud to the same XY extent of the `fvg0` object
# Plot empty voxels between ground and (Zmax - Zcut)
fvg1 <- fvg0 %>% dplyr::filter(emptyVox == 1) # the subset of empty voxels
rgl::open3d()
for(i in 1:nrow(fvg1)) {
  plotVoxel(fvg1$xc[i], fvg1$yc[i], fvg1$zc[i], vx, vy, vz,
  'steelblue', alpha=.3)
}
rgl::axes3d()

rgl::points3d(tls.filtered$X,
        tls.filtered$Y,
        tls.filtered$Z, size = 1)
```

*Box S1: code used to produce figure 2.*
